# A zebrafish embryo screen utilizing gastrulation for identification of anti-metastasis drugs

**DOI:** 10.1101/2022.05.07.490997

**Authors:** Joji Nakayama, Hideki Makinoshima, Zhiyuan Gong

**Affiliations:** Tsuruoka Metabolomics Laboratory, National Cancer Center, Tsuruoka, Japan; Shonai Regional Industry Promotion Center, Tsuruoka, Japan; Department of Biological Science, National University of Singapore, Singapore; Division of Translational Research, Exploratory Oncology Research, and Clinical Trial Center, National Cancer Center, Kashiwa, Japan

**Keywords:** Metastasis, gastrulation, EMT, Phenotyping screening, zebrafish

## Abstract

Few models exist that allow for rapid and effective screening of anti-metastasis drugs. Here, we present a phenotype-based chemical screen utilizing gastrulation of zebrafish embryos for identification of anti-metastasis drugs. Based on the evidence that metastasis proceeds through utilizing the molecular mechanisms of gastrulation, we hypothesize that chemicals which interrupt zebrafish gastrulation might suppress metastasis of cancer cells. Thus, we developed a drug screening protocol which uses epiboly, the first morphogenetic movement in gastrulation, as a marker. The screen only needs zebrafish embryos and enables hundreds of chemicals to be tested in five hours through observing epiboly progression of a test chemical-treated embryos. In the screen, embryos at the two-cell stage are firstly corrected and then developed to the sphere stage. The embryos are treated with a test chemical and incubated in the presence of the chemical until vehicle-treated embryos develop to 90% epiboly stage. Finally, positive ‘hit’ chemicals that interrupt epiboly progression are selected through comparing epiboly progression of the chemical-treated embryos with that of vehicle-treated embryos under a stereoscopic microscope. Previous study subjected 1,280 FDA-approved drugs to the screen and identified Adrenosterone and Pizotifen as epiboly-interrupting drugs. These drugs were validated to suppress metastasis of breast cancer cells in mice models of metastasis. Furthermore, 11β–Hydroxysteroid Dehydrogenase 1 (HSD11β1) and serotonin receptor 2C (HTR2C), which are primary target of Adrenosterone and Pizotifen respectively, promotes metastasis through induction of epithelial-mesenchymal transition (EMT). That indicates the screen could be diverted to a chemical genetic screening platform for identification of metastasis-promoting genes.

## Introduction

Cancer research using zebrafish as a model has attracted attention because this model offers many unique advantages that are not readily provided by other animal models. Futhermore, the zebrafish system has been increasingly recognized as a platform for chemical screening because it provides the advantage of high-throughput screening in an in vivo vertebrate setting with physiologic relevance to humans^1–5^.

Metastasis is responsible for approximately 90% of cancer-associated mortality. It proceeds through multiple steps: invasion, intravasation, survival in the circulatory system, extravasation, colonization, and metastatic tumor formation in secondary organs with angiogenesis^6–8^. Dissemination of cancer cells is an initial step of metastasis and its molecular mechanism involves local breakdown of basement membrane, loss of cell polarity, and induction of EMT^9,10^ These cellular and biological phenomena are also observed during vertebrate gastrulation in that evolutionarily conserved morphogenetic movements of epiboly, internalization, convergence, and extension cooperate to generate germ layers and sculpt the body plan^11^. In zebrafish, the first morphogenetic movement, epiboly, is initiated at approximately four hours post-fertilization (hpf) to move cells from the animal pole to eventually engulf the entire yolk cell by 10 hpf. These movements are governed by the molecular mechanisms that are induced by temporally and spatially regulated gene expression, and these mechanisms and changes in gene expression are partially observed in metastatic progression^12^

### Development of the protocol

Metastasis proceeds through utilizing the molecular mechanisms of gastrulation. At least fifty common genes were shown to be involved in both gastrulation and metastasis progression (Table 1)^13–17^ The fifty genes are expressed in Xenopus or zebrafish embryos, and genetic inhibition of each of the fifty genes in these embryos interferes with gastrulation progression. Conversely, the same fifty genes are ectopically expressed in metastatic cancer cells and confer metastatic properties on cancer cells, and genetic inhibition of each of the fifty genes suppresses metastasis progression. These evidences led us to hypothesize that chemicals which interfere with zebrafish gastrulation might suppress metastasis progression of cancer cells. Based on the hypothesis, we developed a drug screening protocol which uses epiboly, the first morphogenetic movement in gastrulation, as a marker. This screen measures the suppressor effect of each of test chemicals through observing epiboly progression of the chemical-treated embryos (Fig. 1 and Fig. 2).

**Fig. 1.**
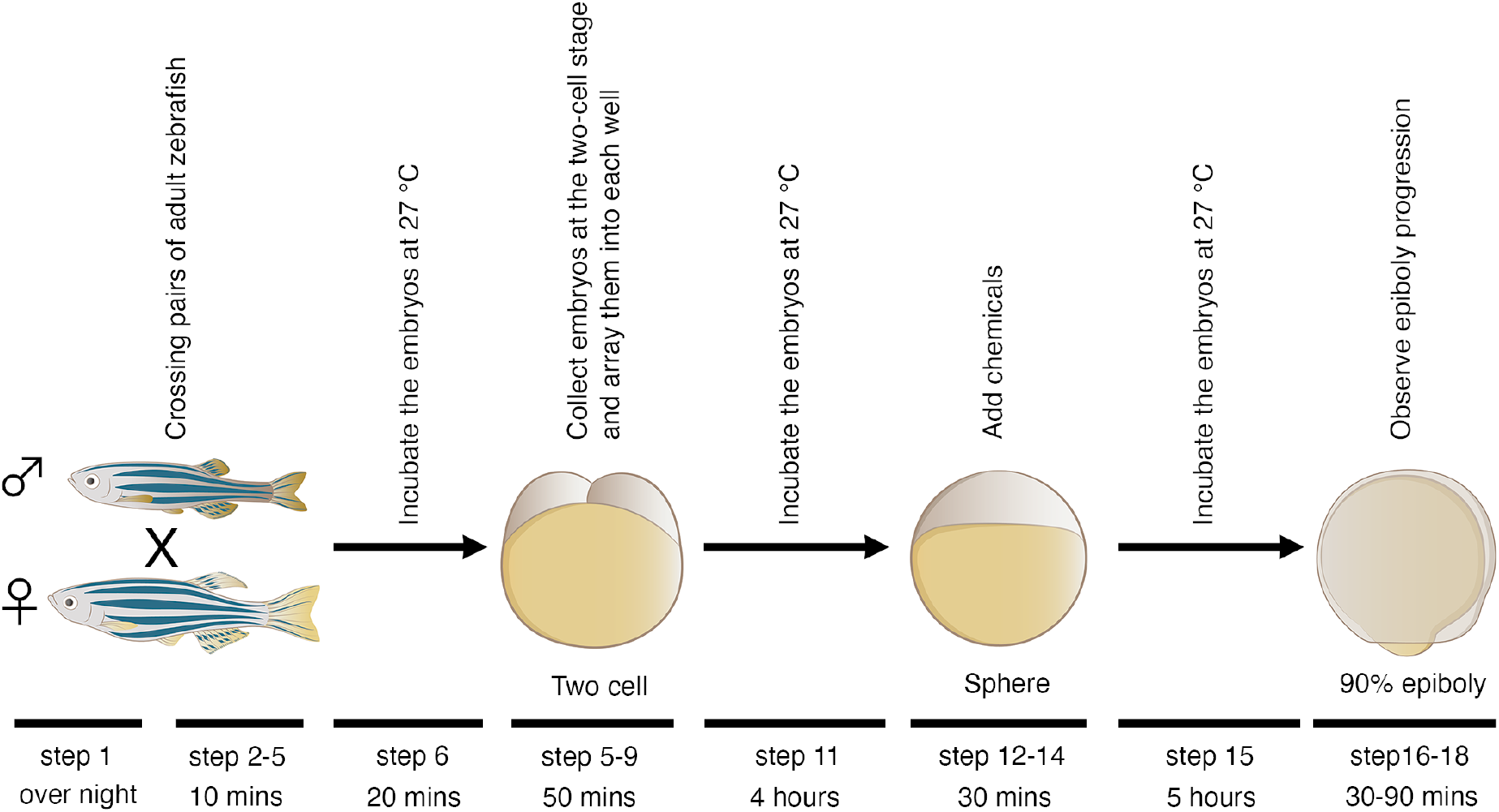
Graphic schematic of a phenotype-based chemical screen using zebrafish embryos. Pairs of adult zebrafish are crossed and their embryos at the two-cell stage are collected and arrayed into individual wells of 24-well plate. Chemicals are added into each well when the embryos develop to the sphere stage. Epiboly progression of each of chemicals-treated embryos are compared with that of DMSO-treated embryos under a stereoscopic microscope when DMSO-treated embryos develop to 90% epiboly stage.

**Fig. 2.**
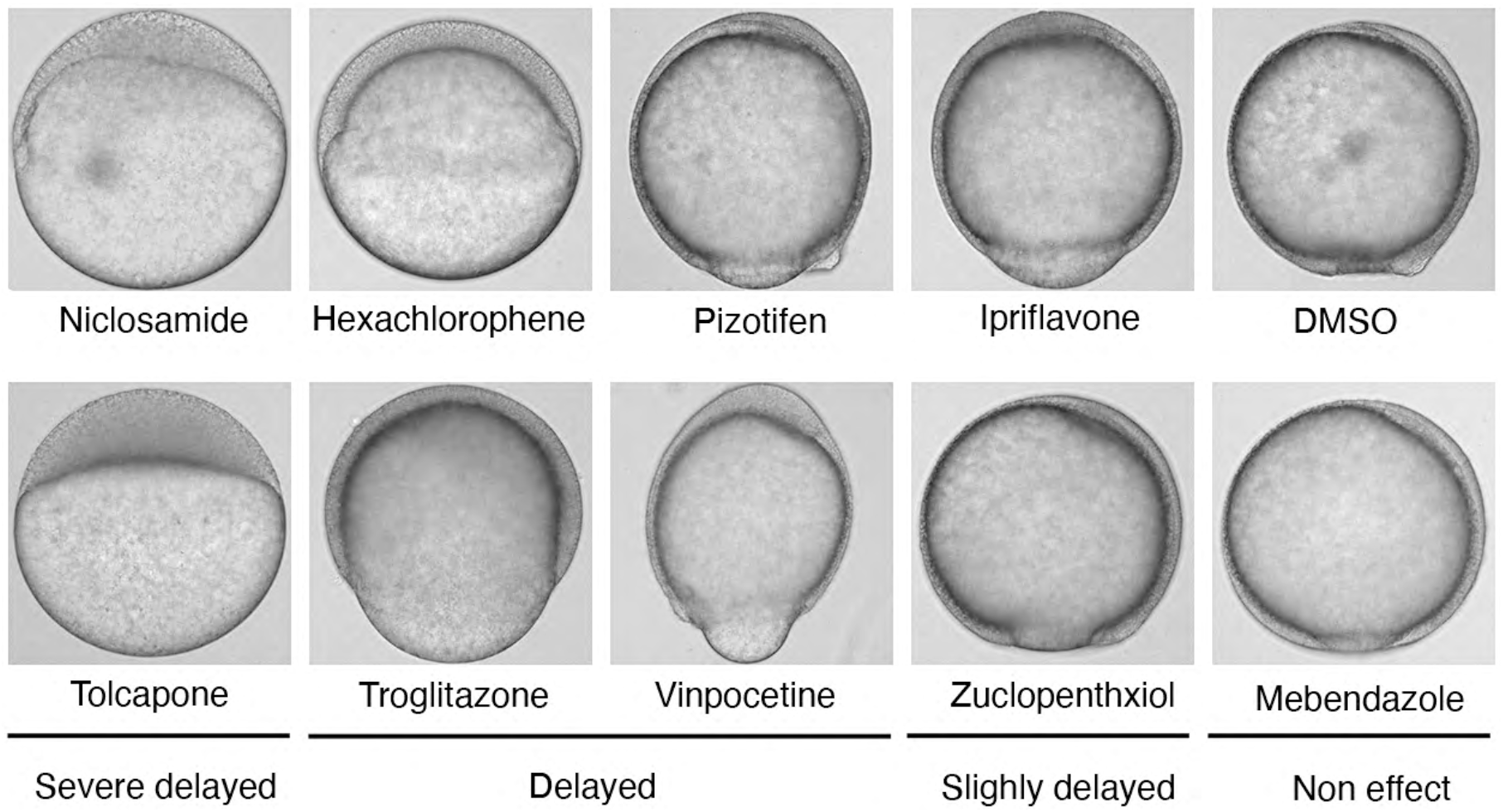
Representative samples of the embryos that were treated with indicated chemicals. Indicated chemicals were added when the embryos develop to the sphere stage. Embryos were treated with 10 μM. Niclosamide-treated embryos serve as positive control and DMSO-treated embryos serve as negative control. Epiboly progression of each of chemicals-treated embryos are compared with that of DMSO-treated embryos under a stereoscopic microscope when DMSO-treated embryos develop to 90% epiboly stage.

**Table 1.**
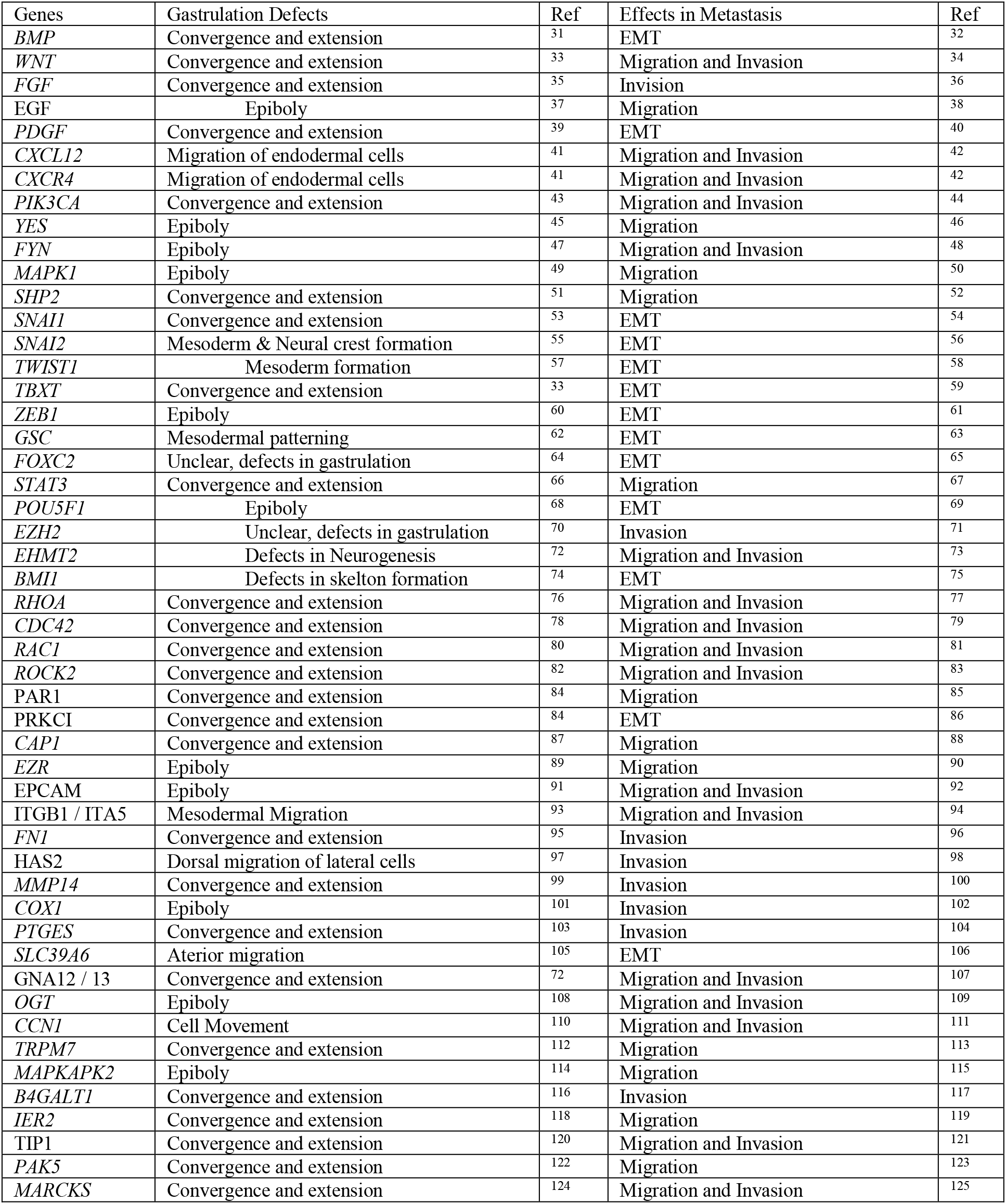
A list of the common fifty genes that are involved between gastrulation and metastasis progression.

### Applications of the method

This screen enables hundreds of chemicals to be tested in five hours. Our study subjected 1280 FDA-approved drugs to this screen and identified Adrenosterone and Pizotifen as epiboly-interfering drugs. These drugs were further validated to suppress metastasis of breast cancer cells in mouse models of metastasis (Fig. 3)^18,19^ This screen can also measure suppressor effect of crude drugs. We subjected 120 herbal medicines to this screen and identified cinnamon bark extract as an epiboly-interfering drug. Cinnamon bark extract was validated to suppress metastatic dissemination of breast cancer cells in zebrafish xenograft model^20^. Moreover, this screen can be diverted to a chemical genetic screening platform for identification of metastasis-promoting genes. HSD11β1 and HTR2C, which are respectively primary targets of Adrenosterone and Pizotifen, induce EMT and promote metastasis of breast cancer cells (Fig. 4)^18,19^.

**Fig. 3.**
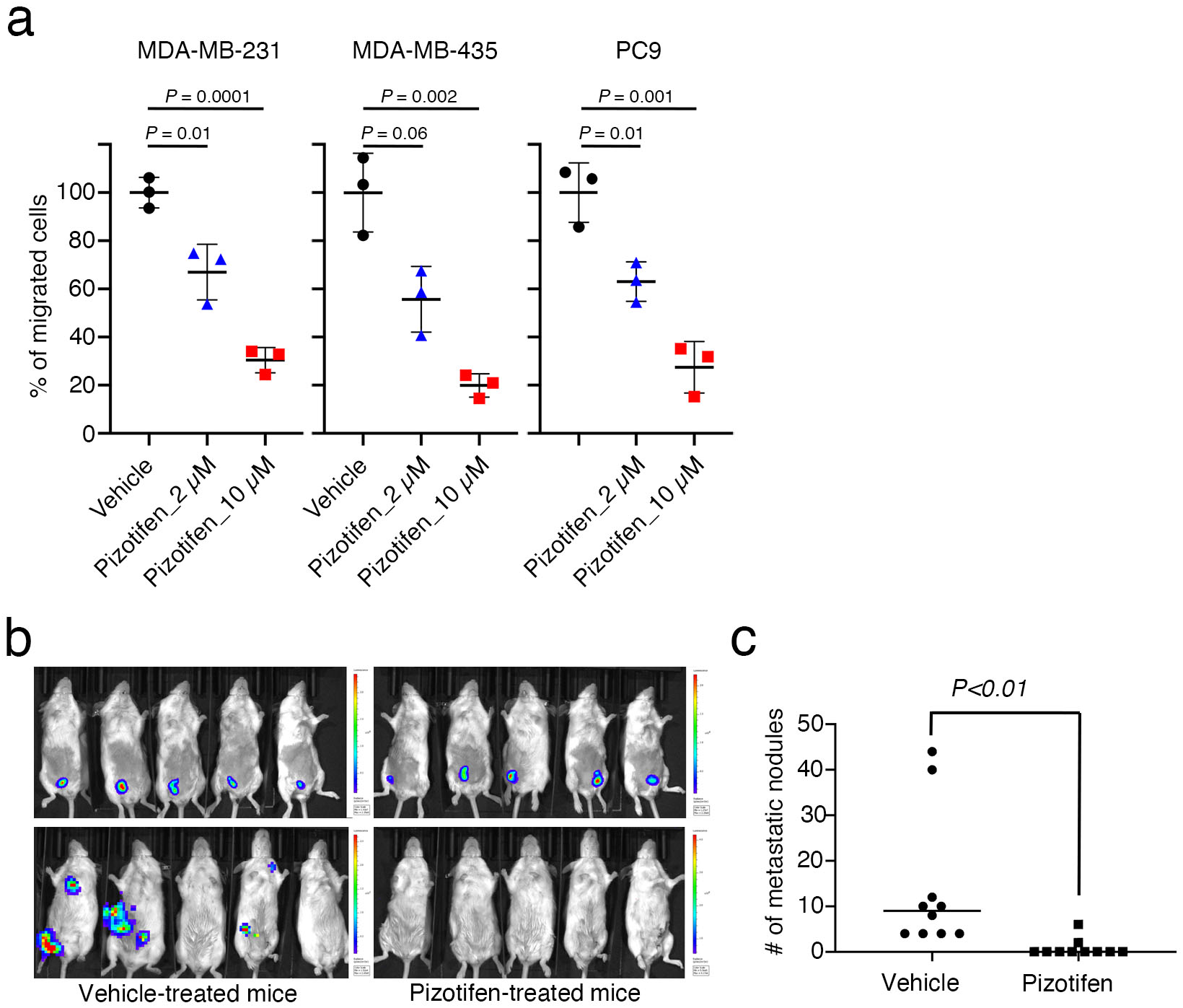
Pizotifen, one of epiboly-interrupting drugs, suppressed metastatic progression of breast cancer cells in vitro and vivo. **a**, Effect of pizotifen on cell motility and invasion of MBA-MB-231, MDA-MB-435, and PC9 cells. Either vehicle or pizotifen treated the cells were subjected to Boyden chamber assays. Fetal bovine serum (1% v/v) was used as the chemoattractant in both assays. Each experiment was performed at least twice. **b**, Representative images of primary tumors on day 10 post-injection (top panels) and metastatic burden on day 70 post-injection (bottom panels) taken using an IVIS Imaging System. **c**, Number of metastatic nodules in the lung of either vehicle-or pizotifen-treated mice.

**Fig. 4.**
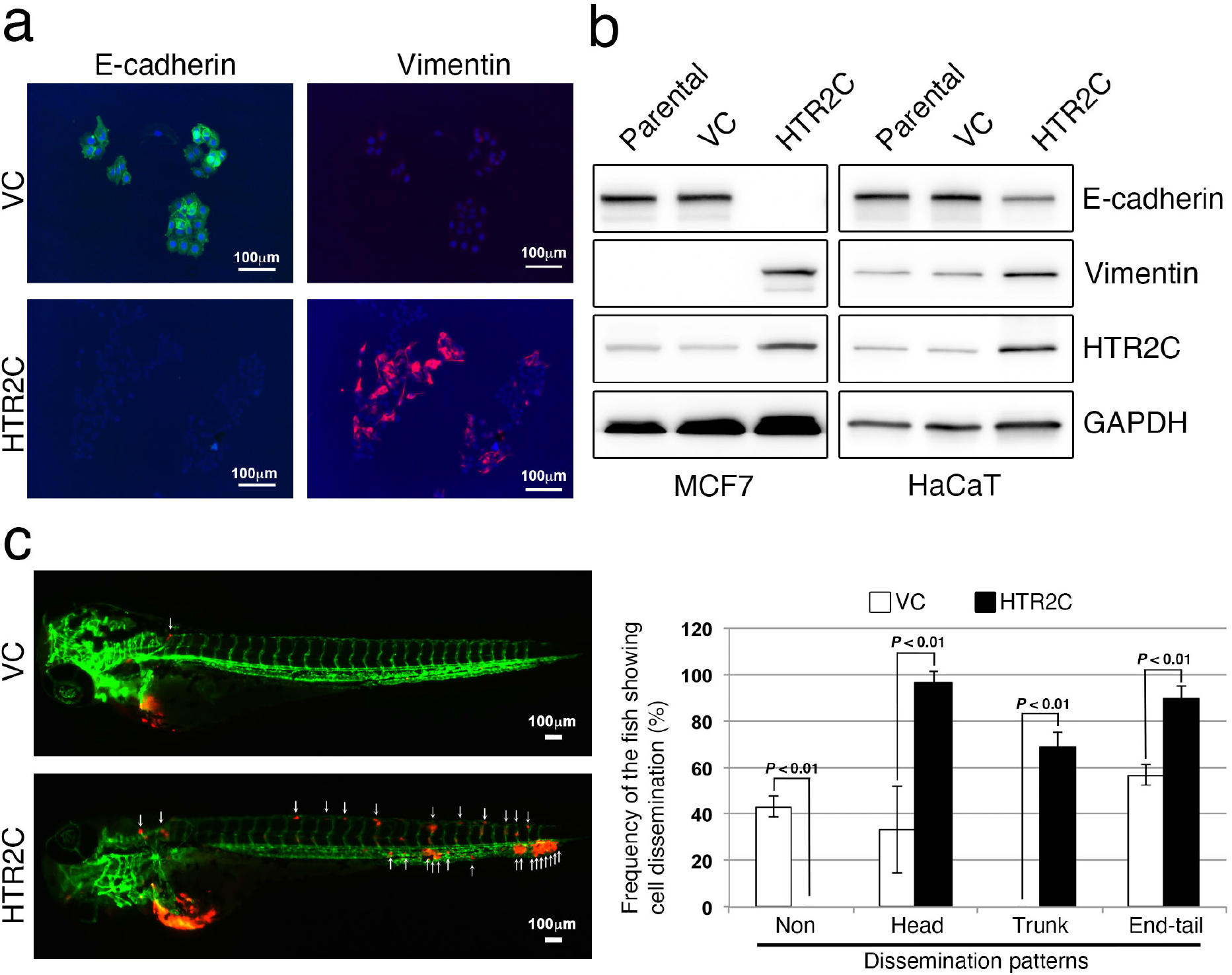
HTR2C, a primary target of Pizotifen, induces epithelial-to-mesenchymal transition (EMT)-mediated metastatic dissemination of human cancer cells. **a**, Immunofluorescence staining of E-cadherin and vimentin expressions in the MCF7 cells. **b**, Expression of E-cadherin and vimentin, and HTR2C were examined by western blotting in the MCF7 and HaCaT cells; GAPDH loading control is shown (bottom). **c**, Representative images of dissemination patterns of MCF7 cells expressing either the control vector (top left) or HTR2C (lower left) in a zebrafish xenotransplantation model. White arrowheads indicate disseminated MCF7 cells. The mean frequencies of the fish showing head, trunk, or end-tail dissemination (right). Each value is indicated as the mean ± SEM of two independent experiments. Statistical analysis was determined by Student’s t test.

### Comparison with other methods

Current mouse models of metastasis are too expensive and time-consuming to use for rapid and high-throughput screening^21,22^ Also in vitro model of metastatic dissemination such as a Boyden chamber assay can test a limited number of chemicals in one assay and needs huge time and effort in analyzing the results^23^. In contrast, our screen only needs zebrafish embryos and enables hundreds of chemicals to be tested in five hours through observing epiboly progression of a test chemical-treated embryos. Furthermore, out of the 78 chemicals which interrupt epiboly progression of zebrafish embryos, 20 of the chemicals were validated to suppress cell motility and invasion of highly metastatic human cancer cells without affecting cell viability in a Boyden chamber assay. Among the 20 chemicals, Adrenosterone and Pizotifen were validated to suppress metastasis of breast cancer cells in mice models of metastasis^18,19^. A disadvantage of this screen is that zebrafish have orthologues to 86% of 1318 human drug targets^27^. Therefore, 75% of the chemicals which interrupt epiboly progression of zebrafish embryos, fail to suppress cell motility and invasion of highly metastatic human cancer cells in a Boyden chamber assay^19^.

### Experimental design

This screen measures suppressor effect of each of chemicals based on epiboly progression of zebrafish embryos. Niclosamid or DMSO is used as positive or negative control, respectively. Epiboly progression of each of chemical treated embryos is compared with that of DMSO-treated embryos. Firstly, embryos at the two-cell stage are firstly corrected and then developed to the sphere stage. The embryos are treated with a test chemical and incubated in the presence of the chemical until vehicle-treated embryos develop to 90% epiboly stage. Finally, positive ‘hit’ chemicals that interrupt epiboly progression are selected through comparing epiboly progression of the chemical-treated embryos with that of vehicle-treated embryos under a stereoscopic microscope^19^.

### Limitations

There is a limitation in delivering chemicals to zebrafish embryos. Zebrafish embryos are surrounded by the acellular chorion, which is known to be about 1.5–2.5 μm thick and to consist of three layers pierced by pore canals. The pore allows passage of water, ions, and chemicals. A study reported molecules which are larger than 3-4 KDa fail to pass through the chorion.

Therefore, this screen may not be able to measure suppressor effect of the molecules which are larger than 3-4 KDa^28^.

## Materials

REAGENTS

- Wild-type zebrafish strain
- E3 medium (5.0 mM NaCl, 0.17 mM KCl, 0.33 mM MgSO_4_)
- FDA, EMA, and other agencies-approved chemical libraries were purchased from Prestwick Chemical (Illkirch, France).
- Niclosamid
- DMSO

EQUIPMENT

- 24-well flat bottom plastic plates (Corning)
- Stereomicroscope (MZ75, Leica)
- Incubator (Thermo)

### Procedure

Zebrafish mating setup (Day 0) _Timing 10 mins

1. On the night before collecting embryos, arrange male and female zebrafish in pairs separated by a divider CRITICAL STEP Young adult zebrafish should be used for the crossing. Qualities of zebrafish embryos affect screening efficiency.

Embryo collection, and distribution (Day 1) _Timing 10-30 min

2. Remove the divider to allow the fish to spawn.
3. To obtain zebrafish embryos of the same development stage, zebrafish were crossed for 10 mins. If more than twenty chemicals were tested, the crossing were conducted three times at three different time points (Group A_8:30, Group B_9:00 and Group C_9:30AM).
4. After 10 mins, set back divider to prevent zebrafish from spawning. CRITICAL STEP This screen measures suppressor effect of each chemical on progression of epiboly in live zebrafish embryos. Therefore, epiboly proceeds during measuring the effect under a stereoscopic microscope. If more than 20 chemicals are tested, screening should be divided into more than two sessions and each of the sessions start at different time point. For example, if 60 chemicals are screened, zebrafish should be crossed at three different time points over 30 mins apart. To do that, 30 mins for measuring the effects would be ensured.
5. Collect the embryos and remove dead embryos
6. Incubate the embryos at 27 °C for twenty mins
7. Collect embryos at the two-cell stage under stereoscopic microscope.
8. Array approximately twenty embryos into each well of a 24-well plate
9. Remove E3 medium from each well including the embryos by using a pipet
10. Add 900 μl of E3 medium to the well.

Embryo development to the sphere stage_Timing 4 hours

11. Incubate the embryos at 27 °C until the embryos develop to the sphere stage CRITICAL STEP The temperature of E3 medium affects the rate of development of zebrafish embryos. Higher temperature accelerates the rate; conversely, lower temperature slows the rate^29^ Therefore, non-uniform temperature between E3 medium of each well of 24-well plate containing zebrafish embryos would cause false positive.

### Addition of chemicals_Timing 30 mins

12. At 30 mins before adding test chemicals to embryos, prepare 10-fold concentration of each of the chemicals in E3 medium
13. Add 100 μl of 10-fold concentration of the medium to 900 μl of E3 medium containing zebrafish embryos when the embryos develop to the sphere stage.
14. For example, for 60 test chemicals to be screened, they are divided into three groups.
  a. First 20 test chemicals plus Niclosamid as positive control, and DMSO as negative control are added into group A when embryos from group A develop to sphere stage.
  b. Second 20 test chemicals plus Niclosamid, and DMSO were added into group B when embryos from group B develop to sphere stage.
  c. Last 20 test chemicals plus Niclosamid, and DMSO were added into group C when embryos from group C develop to sphere stage

### Development of DMSO-treated embryos to 90% epiboly stage_Timing 5 hours

15. After the addition of test chemicals, the embryos are incubated at 27 °C for approximately five hours. CRITICAL STEP The temperature of E3 medium affects the rate of development of zebrafish embryos. Non-uniform temperature between E3 medium of each well of 24-well plate containing zebrafish embryos would cause false positive.

### Measuring the inhibition effects of each of chemicals_Timing 30 mins

16. Comparing epiboly progression of each of chemicals-treated embryos from group A with that of DMSO-treated embryos from group A under the stereoscopic microscope when DMSO-treated embryos from group A develop to 90% epiboly stage.
17. After approximately 30 min from observing embryos in group A, measuring the inhibition effects of each of chemicals on epiboly progression of embryos in group B and when DMSO-treated embryos from group B develop to 90% epiboly stage.
18. After approximately 30 mis from observing embryos in group B, measuring the inhibition effects of each of chemicals on epiboly progression of embryos in group C and when DMSO-treated embryos in group C develop to 90% epiboly stage. CRITICAL STEP Epiboly proceeds during comparing epiboly progression of each of chemicals-treated embryos with that of DMSO-treated embryos under the stereoscopic microscope. Therefore, measuring the effect should be done in 30 mins.

### Timing

Step 1, Zebrafish mating setup: overnight

Step 2-5, Crossing zebrafish: 10 mins

Step 6, Develop the embryos to the two-cell stage

Step 7, Collect two cell stage embryos: 20 mins

Step 8-10, Array 20 embryos into each well of 24-well plate: 30 mins

Step 11, Develop the embryos to sphere stage: 4 hours

Step 12-14, Prepare chemical drugs in E3 medium and add the medium into each well of 24-well plate: 30 mins

Step 15, Develop the embryos to 90% epiboly stage: 5 hours

Step 16-18, measuring epiboly progression: 30 mins

### Troubleshooting

Qualities of zebrafish embryos affect screening efficiency. Low qualities of embryos show high frequencies of abnormal embryos with asymmetric cell cleavage, and development of the embryos arrest at early cleavage stages^30^. If a screen used low qualities of zebrafish embryos, it would generate false ‘hit’ chemicals since suppressor effect of a test chemical is measured by observing epiboly progression of the chemical-treated embryos. If the number of zebrafish embryos showing morphological abnormalities correlate with final concentration of a test chemical, the abnormalities may result from an effect of the test chemical on the embryos.

### Anticipated results

Suppressor effects of a tested chemical on epiboly progression of zebrafish embryos are significantly affected by final concentration of the chemical. Previous study subjected 1,280 FDA-approved drugs to the screen and showed 6% (78/1280) of the tested drugs affected epiboly progression of the embryos when the embryos were treated with 10 μM. Out of the 78 epiboly-interrupting drugs, 25% of the drugs succeed to suppress cell motility and invasion of highly metastatic human cancer cells in a Boyden chamber assay. In contrast, epiboly progression was affected more severely when the embryos were treated at 50 μM. 10.3% (132/1280) of the tested drugs affected epiboly progression of the embryos, but 85 % (112/132) of the epiboly-interrupting drugs failed to suppress cell motility and invasion of highly metastatic human cancer cells in a Boyden chamber assay^19^.

## Author contributions statements

J.N. created a concept of this screen, designed research, conducted experiments, analyzed data, wrote an original draft, supervised this project; H.M. and Z.G. administered this project, acquired funding. All authors reviewed and edited the draft.

## Acknowledgments

We thank Dr. Herrick (Albany Medical College) for providing pCMV-h5TH2C-VSV with us. Illustrations of Fig. 1 were drawn by Ami Inoue (Kyoto University of the Arts). This study was funded by grants from National Medical Research Council of Singapore (R-154000547511) and Ministry of Education of Singapore (R-154000A23112 and R154000B88112) to ZG. Experiments using mice were supported in part by research funds from the Yamagata prefectural government and the City of Tsuruoka.

